# Single cell atlas of beige remodeling of white adipose tissue reveals a myeloid to lymphoid shift during cold exposure compared to beta 3 adrenergic stimulation

**DOI:** 10.1101/2020.05.30.125146

**Authors:** Nabil Rabhi, Anna C. Belkina, Kathleen Desevin, Briana Noel Cortez, Stephen R. Farmer

## Abstract

White adipose tissue (WAT) is a dynamic tissue, which responds to environmental stimuli and dietary cues by changing its morphology and metabolic capacity. The ability of WAT to undergo a beige remodeling has become an appealing strategy to combat obesity and its related metabolic complications. Within the cell mixture that constitutes the stromal vascular fraction (SVF), WAT beiging is initiated through expansion and differentiation of adipocytes progenitor cells, however, the extent of the SVF cellular changes is still poorly understood. Additionally, direct beta 3 adrenergic receptor (Adrb3) stimulation has been extensively used to mimic physiological cold-induced beiging, yet it is still unknown whether Adrb3 activation induces the same WAT remodeling as cold exposure. Here, by using single cell RNA sequencing, we provide a comprehensive atlas of the cellular dynamics during beige remodeling within white adipose tissue. We reveal drastic changes both in the overall cellular composition and transcriptional states of individual cell subtypes between Adrb3- and cold-induced beiging. Moreover, we demonstrate that cold exposure induces a myeloid to lymphoid shift of the immune compartment compared to Adrb3 activation. Further analysis, showed that Adrb3 stimulation leads to activation of the interferon/Stat1 pathways favoring infiltration of myeloid immune cells, while repression of this pathway by cold promotes lymphoid immune cells recruitment. These findings provide new insight into the cellular dynamics during WAT beige remodeling and could ultimately lead to novel strategies to identify translationally-relevant drug targets to counteract obesity and T2D.

## INTRODUCTION

Adipose tissue is a central metabolic organ for whole body energy homeostasis. An imbalance between energy intake and energy expenditure increases adiposity and can lead to severe metabolic disease (Sun et al., 2011). White adipose tissue (WAT) plays a key role as a reservoir for triglyceride storage whereas brown adipose tissue (BAT) dissipates energy as heat through mitochondrial uncoupling. Under appropriate stimulation, such as cold exposure or beta-3- adrenergic receptor (ADRB3) stimulation, WAT can adopt a thermogenic phenotype, sustained by emergence of uncoupling protein 1 (UCP1) expressing cells (Kajimura et al., 2015). These cells, called beige or brite fat cells, share the same energy-burning capacity as BAT through substrate oxidation, but present a distinct molecular and developmental origin. Increasing whole body thermogenic capacity by activating BAT and promoting beige cells emergence may represent a promising strategy to counteract the development of obesity and diabetes (Cannon and Nedergaard, 2004, Bartelt et al., 2011). Indeed, activation of thermogenesis plays a critical role in promoting a shift in energy expenditure in obese and T2D individuals through a potent glucose and lipid clearance to fuel thermogenesis. Essentially all studies to date show that beiging of WAT prevents high-fat, diet-induced insulin resistance and weight gain, resulting in positive metabolic indicators, such as insulin sensitivity and euglycemia (Berbée et al., 2015). Additionally, adult human BAT has been identified with more of a brite/beige character than classic rodent brown fat (Sharp et al., 2012, Cypess et al., 2009). Therefore, a better understanding of mechanisms controlling beige adipogenesis could lead to the development of new therapies for metabolic diseases.

WAT is a complex organ consisting of a mixture of mature adipocytes and stromal vascular cells (SVC). SVCs, comprising 80% of WAT cells, are a dynamic and complex assortment of resident immune cells, vascular cells, mesenchymal stem cells (MSC) and pre-adipocytes that can change with development and WAT remodeling (Eto et al., 2009). Furthermore, these changes in cell population can play an important role in the capacity of the tissue to respond to the metabolic needs of the body (Kahn et al., 2019, Choe et al., 2016). While a lot of effort has been devoted to defining the cellular plasticity during obesity, little is known about the landscape of these changes during early beige adipogenesis. Although some studies have attempted to define beige adipogenesis, the focus has been on characterizing progenitor cells origin and fate decisions using either mouse lineage tracing models or cells sorting (Rajbhandari et al., 2019, Vishvanath et al., 2016, Sanchez-Gurmaches and Guertin, 2014). Moreover, previous investigations have used an ADRB3 activator (CL316,243) as the tool to model WAT response to cold (Lee et al., 2017, Burl et al., 2018, Rajbhandari et al., 2019). Indeed, ADRB3 activation in vivo by CL 316,243 (CL) provides a means to rapidly induce WAT beiging, however, whether CL induces the same adipose tissue remodeling as cold exposure remains to be determined. Herein, we provide a comprehensive atlas of WAT SVC cellular subtypes and address the change in complexity of the tissue during early response to cold and CL using single cell RNA sequencing (scRNAseq) technology. Our results identify critical cell subpopulations and their dynamic changes that occur following cold or CL treatment. In combination with flow cytometry, we demonstrate that immune cells with a myeloid origin expand in response to CL treatment while cold exposure leads to expansion of lymphoid cells mainly B cells, CD4 and CD8 T cells. Mechanistically, this immune shift is controlled by activation of the interferon pathway and Stat-1 phosphorylation.

## RESULTS

Histological analysis of C57BL/6J mice treated with Adrb3 agonist (CL316,243; CL) or cold for either 3 days or 10 days led to a comparable level of beiging within the subcutaneous inguinal adipose tissue (iWAT) (**Figure S1A**). However, gene expression analysis of immune markers within the SVC fraction showed a decrease of Ccl2; the monocytes/ macrophages chemoattractant cytokine in cold conditions while no changes were observed in response to CL at both 3 and 10 days (**Figure 1A**). In agreement with these results, we found that Adrge1, a macrophage marker was increased in CL treated mice while decreased by cold exposure at both monitored time points (**Figure 1A**). Moreover, extracellular matrix (ECM) markers such as Col1a1 and Col3a1 were increased in response to CL and decreased following cold exposure. Interestingly, adipogenesis markers Fabp4 and adiponectin were only increased by cold exposure at 3 days (**Figure 2A**) and thermogenic markers were more responsive to CL treatment than cold. To gain more insight into the differences between cold- and Adrb3- induced immune response, we stained for CD45, a common immune cell marker (**Figure 1B**). We found that CL increases immune cells infiltration at both 3 and 10 days whereas no changes were observed with cold exposure (**Figures 1B** and **1C**). Surprisingly, we found that more T cells were observed following cold exposure than CL treatment at both monitored times (**Figures 1D and 1E**). Although both CL and cold induce beiging, our data suggest that the mechanisms leading to the beige phenotype have an immune component.

**Figure 1:**
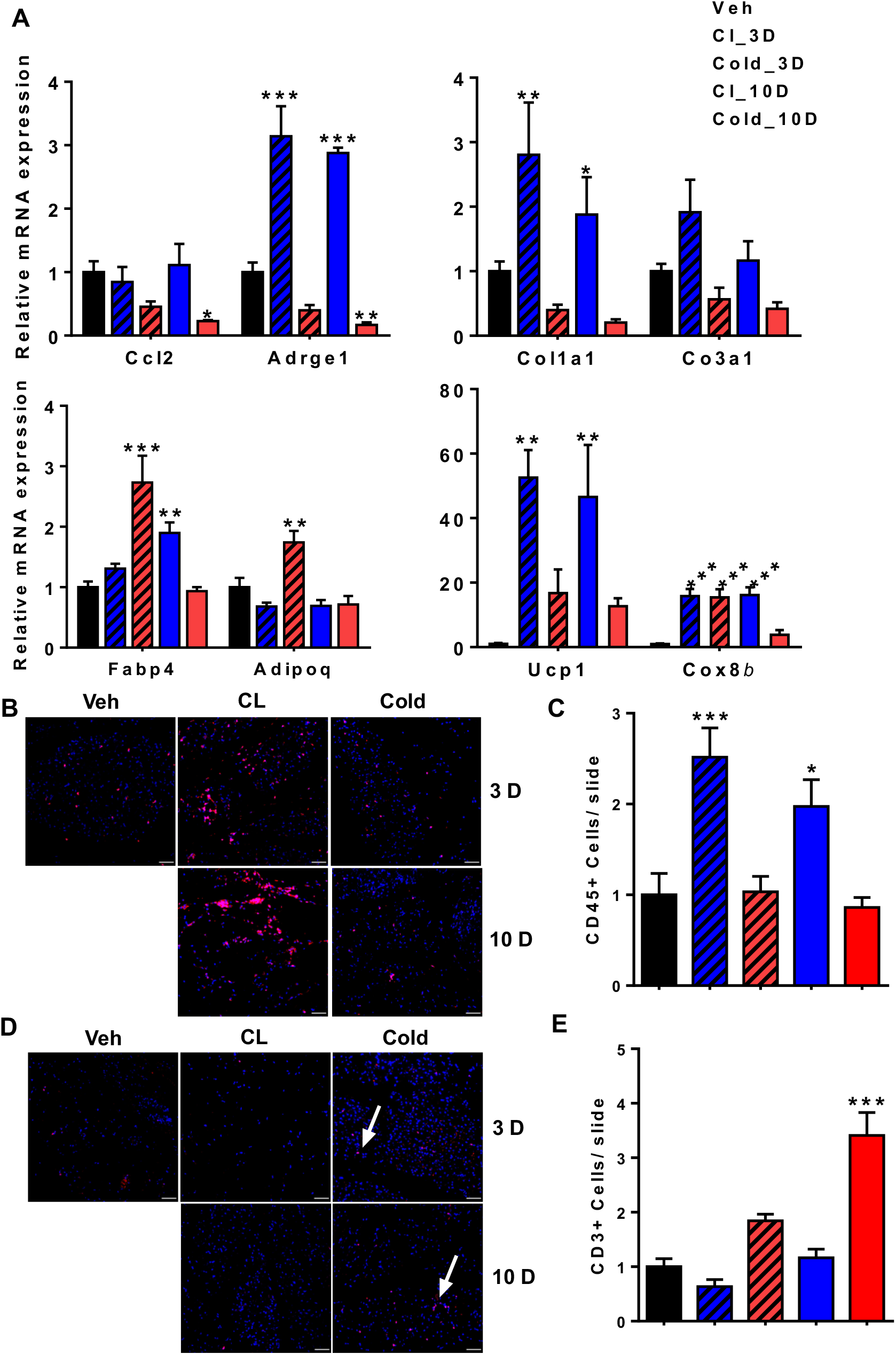
CL and cold treatment lead to distinct beige remodeling. All experiments were carried out following 3 days of in vivo CL316,234 or cold exposure. (A) Real time PCR analysis of relevant immune (top left), ECM (top right), adipogenesis (bottom left) and thermogenesis markers (bottom right) in the SVF from iWAT. (B) Representative image of CD45 immunofluorescent staining. (C) Quantification of CD45+ cells (n=5) (D) Representative image of CD3 immunofluorescent staining. (E) Quantification of CD453+ cells (n=5). Data are presented as mean ± SEM. p-values. n=5-6 animals in each group; *p < 0.05, **p < 0.01, ***p < 0.001.

**Figure 2:**
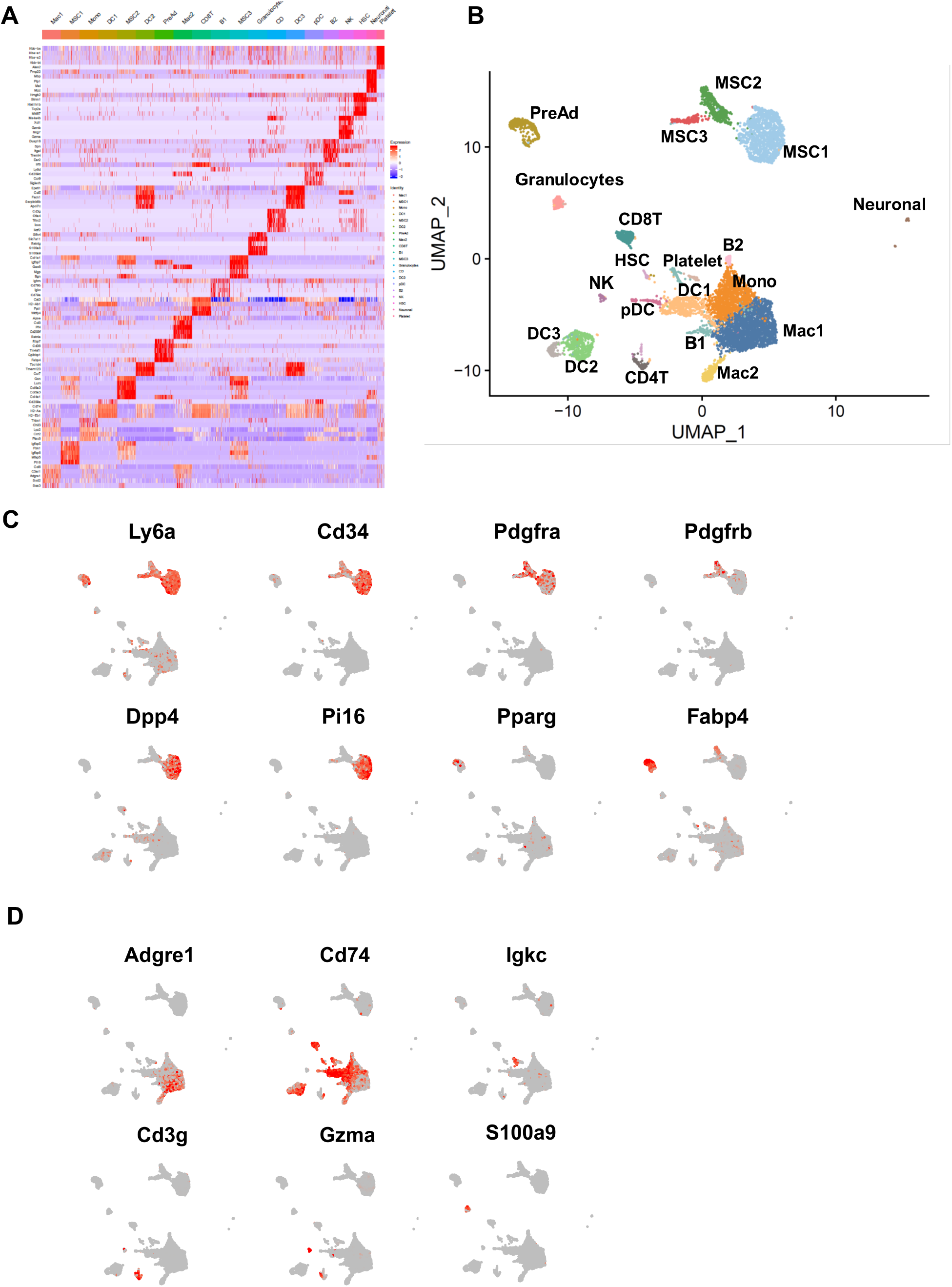
Single cell sequencing reveals cellular heterogeneity of the SVF from iWAT. (A) Gene-expression heatmap showing the top 5 most differentially expressed (DE) genes (ordered by) used for biological identification of each cluster compared to all other clusters. Genes are represented in rows and ordered by decreasing P value and cell clusters in columns. (B) UMAP plots of iWAT SVF populations from combined dataset of control, CL and cold treated mice for 3 days representation, which identified 20 clusters from 6022 cells. UMAP expression plots of representative DE genes displaying expression of known as in (C) adipocytes progenitors’ cells markers and in (D) immune cells markers overlaid in red across all populations. Scale bars represent z-test–normalized gene expression in A and gene counts in C and D. MSC indicates mesenchymal stem cells; PreAd, committed pre-adipocytes; Mac1, M1 macrophages; Mac2, M2 macrophages; Mono, monocytes; DC, dendritic cells; pDC, Plasmacytoid dendritic cell; CD4T; CD4+ T cells, CD8T, CD8+ T cells; B, B cells; NK, natural killer cells; HSC, myeloid progenitor cells.

To understand the full extent of the divergent cellular response to CL and cold, we performed scRNA-seq analysis of the SVC fraction isolated from iWAT of control, CL or cold treated mice for 3 days to capture early changes. Data from all three treatments were pooled together to identify subpopulations within the captured cells. Unsupervised clustering using gene markers singled out 20 distinct cell clusters (**Figure 2A**). We assigned putative biological identities to each cluster by manual annotation using established gene expression patterns as well as by interrogating a gene expression atlas (Su et al., 2004, Ravasi et al., 2010). The annotation resulted in several groups of cells including pre-adipocytes, mesenchymal stem cells, immune cells and neuronal cells (Figure 2B). Previous reports identified two to three cellular subpopulations with an adipogenic potential that were defined as mesenchymal progenitor cells (MSC) (Merrick et al., 2019, Hepler et al., 2018, Burl et al., 2018). However, these data were either generated from Pdrgfrb+ sorted cells or assuming that canonical MSC markers Cd34, Pdgfra, Lys6a (Sca1) are co-expressed within the same cell. Because CD34 and Sca1 are the major markers of stemness for most of mice progenitors, we plotted them across our single cell data to discriminate between possible MSC and other cells subtype. We found that four clusters which we named MSC1, MSC2, MSC3, and pre-adipocytes express both markers (**Figure 2B** and **2C**). Pdgfra was expressed in most of the cells in the three MSC subpopulations and was highly expressed in MSC2 cluster. Dpp4 and Pi16, markers previously proposed as interstitial progenitors, were exclusively expressed in MSC1. Fabp4 was expressed in both MSC2 and preadipocytes and Ppary was exclusively expressed by the preadipocytes cluster suggesting that the MSC2 state may precede the pre-adipocytes state (**Figure 2C**). Genes encoding ECM components were expressed at different levels within the four clusters with MSC2 cluster expressing the largest number of ECM related genes (**Figure 2A**, **Table S1**). Interestingly, collagen types expression was found to be different between populations. Indeed, Col14a was identified as a marker for MSC1 cluster, Col15a as a marker for MSC2 cluster while both marked MSC3 cluster (**Table S1**).

Our merged data from the 3 conditions allowed for the identification of 13 distinct immune clusters (**Figure 2B, Table S1**). We used unsupervised annotation and cell type-specific markers to interpret and identify the resulting 13 immune clusters based on literature searches and the Immunological Genome project database (**Figure 2D**)(Jojic et al., 2013, Yoshida et al., 2019). We identified 3 clusters that express macrophages markers such as Adgre1 (**Figure2**D). Proinflammatory cytokines such Ccl2, Ccl6, Ccl9, Ccl12, Cxcl2 were highly expressed in M1 macrophages cluster (Mac1); Mrc1, Cd 209, Lyve1, Cd36 and Mmp9 expressing macrophages were annotated as M2 macrophages (Mac2) and a mixed monocytes/macrophages cluster was marked by the expression of Ccr2, Lyz2, Cy6c2, Ms4a4c, Tyrobp and Cd52 (**Figure 2A, Table S1**). Four clusters were annotated as dendritic cells (DC1, DC2, DC3 and pDC) expressing the common DC cells markers such as CD74 and two B cells clusters expressing genes such as Igkc (**Figure 2A and 2D, Table S1**). We also identified Cd4-T-cells (CD4T), Cd8-T-cells (CD8T), natural killer cells (NK) and a mixed population of granulocytes (**Figure 2A, 2B and 2D, Table S1**). In conclusion, our scRNAseq revealed twenty subpopulations of cells with distinct markers, although the functions of these subsets remain to be elucidated. These findings provide a point of reference for examining occurrences in each cell subset during beiging of adipose tissue.

To commence such an examination, we performed a side-by-side comparison of the control, CL and cold treated datasets. While the data showed that the twenty clusters are represented within the three conditions, our analysis revealed drastic changes both in the overall cellular composition and transcriptional states of individual cell subtypes (**Figure 3A** and **3B**). Initial analysis of Uniform Manifold Approximation and Projection (UMAP) maps of the data showed that the pre-adipocytes population increased in CL condition compared to cold and control treatment (Figure 3A). However, normalization of the data to total number of cells sequenced per condition revealed that only cold increased the pre-adipocytes population (**Figures 3B** and **S2A**). MSC3 cluster showed the same pattern while both MSC1 and MSC2 cluster were reduced by both cold and CL. However, the reduction was more prominent with CL than cold. (**Figure 3B**). Normalized macrophages and monocytes populations were increased by CL and reduced by cold. In contrast, CD4T cells NK and all B cells were increased by cold and reduced by CL treatment (**Figure 3B**). These results suggest a dissimilar immune cell response to CL and cold. Interestingly, the cell populations that are increased with CL are mostly from a myeloid origin whereas cold promotes an increase of immune cells of lymphoid origin (**Figure 3C**). We verified the results obtained by scRNAseq using flow cytometry of the iWAT SVF from mice treated with vehicle, CL or cold for 3 days. We used a panel of antibodies that allowed detailed assessment of multiple immune subsets previously identified by the scRNAseq including B cells, CD4+ and CD8+ T cells, NK cells, DC cells, granulocytes, M1 and M2 macrophages and monocytes (**Figures S3B** and **S3C**). In concordance with previous results, immune cells with a myeloid origin were reduced by cold compared to veh or CL treated mice (**Figure 3D**). More importantly, cold induced lymphoid origin immune cells including B cells (CD19+), CD4 and CD8 T cells compared to the other treatments. All together these data suggest that cold- and CL-induced beiging involve a different immune remodeling leading to the same level of UCP1+ adipocytes (**Figure S1A**). Indeed, activation of Adrb3 induces a specific activation of cells with a myeloid origin such as macrophages while cold leads to increased recruitment of immune cells with a lymphoid origin suggesting that CL treatment isn’t able to activate the full immune system to mimic the cold.

**Figure 3:**
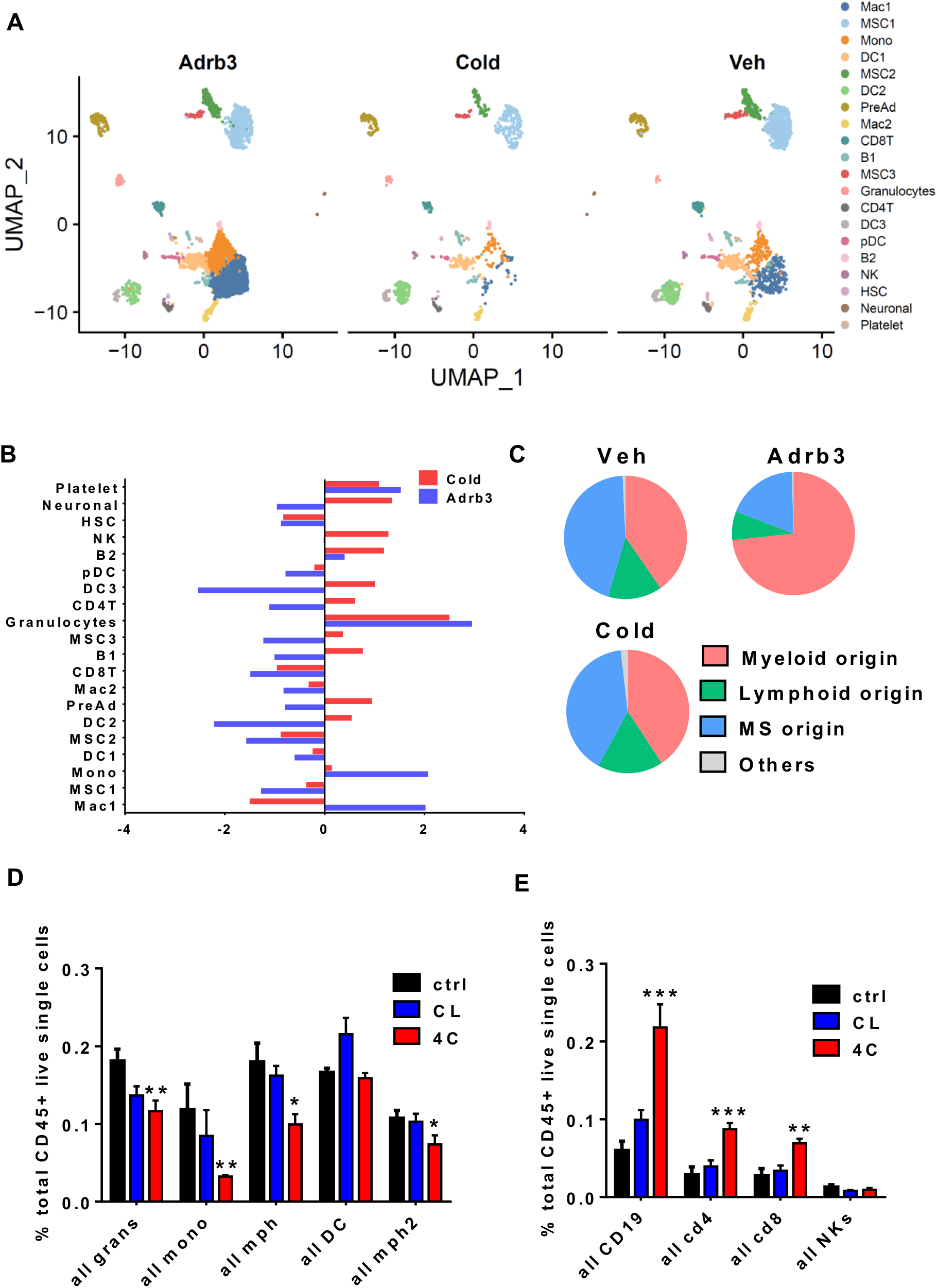
CL and cold differently alter the adipose resident immune compartment. (A) Side- by-side UMAP plots of iWAT SVF populations from control, CL or cold treated mice for 3 days. (B) Bar chart showing population fold-changes in relative abundance of each cluster induced by 3 days of CL or cold treatment compared to vehicle treated mice. (C) Proportions of cell populations grouped by the cellular origin. (D) Quantification of frequency of selected myeloid cell subsets in the iWAT depot after 3 days of CL or cold treatment compared to vehicle treated mice (n=6) performed by flow cytometry. (E) Quantification of frequency of selected lymphoid cell subsets in the iWAT depot after 3 days of CL or cold treatment compared to vehicle treated mice (n=6) performed by flow cytometry. Data are presented as mean ± SEM. p-values. n=5-6 animals in each group; *p < 0.05, **p < 0.01, ***p < 0.001.

To gain more insight into the mechanisms controlling the shift from myeloid to lymphoid immune cells recruitments upon cold exposure compared to CL, we performed differential gene expression analyses between the same clusters using the treatments as a variable. The results showed that both immune cells with a myeloid origin or a lymphoid origin activate a different set of genes in response to either CL or cold (**Figures S4A** and **S4B**). Because the activation of the monocytes and macrophages precedes the activation of myeloid lymphoid cells such as T cells and B cells, we focused on macrophages. Differential analysis revealed that cold induced the up regulation of 140 genes while CL up regulated 23 genes only 9 of those were overlapping with cold. We also identified 57 genes down regulated by cold while CL only decreased 2 genes (**Figures 4A** and **4B**). Gene ontology analysis of genes down regulated by cold showed an enrichment of genes associated with biological processes including cytokine mediated signaling, cellular response to type 1 interferon and type 1 interferon signaling pathway (**Figure 4D**). We next examined the expression of genes induced by interferon such as Irf7, Isg15, Ifit1 and Saa3 across all the identified clusters. Surprisingly, we found that interferon target genes are induced in most immune population regardless of their origin. To further confirm these results, we stained tissue from vehicle, CL or cold treated mice with antibody against pStat1, an interferon induced signaling component. Our data showed that CL treatment induced a considerable phosphorylation of Stat1 in SVF cells. Collectively, these results strongly suggest that repression of the interferon/pStat1 pathway controls the shift from myeloid to lymphoid immune cells recruitment during cold exposure compared to CL.

**Figure 4:**
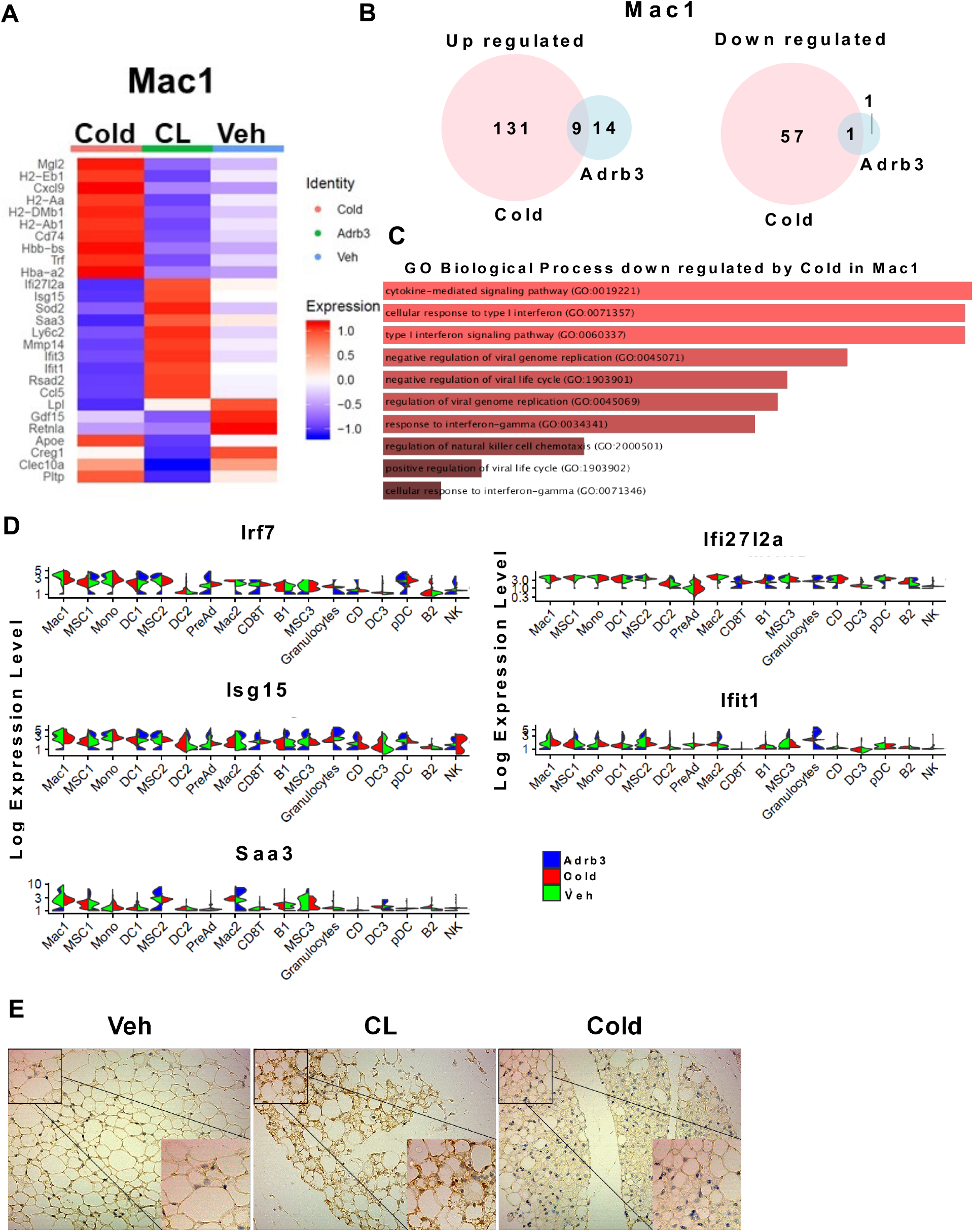
CL-induced beiging leads to an interferon/Stat1 response. (A) Heatmap of differentially expressed genes in Mac1 cluster of iWAT from mice after 3 days of CL or cold treatment. (B) Venn diagram of up (left) and down (right) regulated genes within Mac1 cluster after 3 days of CL or cold treatment (C) GO-driven pathway analysis of DE down regulated gene by cold in Mac1 (D) Violin plots for the expression of interferon response genes (Expression in each cell is shown along with the probability density of gene expression). (E) representative images of immunohistochemistry of P-Stat1 (Tyr701) in iWAT section from mice treated for 3 days with vehicle, CL or cold.

## DISCUSSION

In the present study, we reveal a complete atlas of cellular complexity of the SVF during beige remodeling. All cell populations were present within iWAT from control mice suggesting that cells are at a paused-like state to maintain tissue homeostasis. Cold and CL treatment lead to a large modulation of the cellular composition to achieve a beige phenotype. Previous studies have identified two distinct MSC population within the epidermal WAT (eWAT) and three MSC within the iWAT (Merrick et al., 2019, Burl et al., 2018, Hepler et al., 2018). Our current work reveals the existence of four distinct MSC populations harboring an adipogenic potential which express classical stemness markers and can be distinguished by specific markers including different collagen subtypes. The co-existence of the four populations could explain the high adipogenic potential of the iWAT compared to the eWAT. While all MSCs exist within the control mice iWAT, cold and CL treatments lead to major changes both in the cells number and signaling pathways of individual MSCs subtypes. Interestingly, pre-adipocytes and MSC3 cluster were increased in response to cold compared to CL suggesting that cold leads to significantly more expansion of those populations. However, further studies will be needed to determine if there are any differences between cold and CL stimulation in the recruitment potential to adipocytes progenitors.

Large changes within the immune fraction composition were also observed. In agreement with previous reports from both whole tissue RNA sequencing and scRNAseq, we found that Adrb3- induced beiging leads an increase of macrophages recruitment (Lee et al., 2016, Nguyen et al., 2011, Burl et al., 2018). At a more global scale, immune cells derived from myeloid origin were increased in response to CL compared to cold. In contrast, cold promoted the recruitment of lymphoid originated immune cells including B cells, CD4 and CD8 T cells. This suggests a shift from myeloid to lymphoid immune cells is an important step to promote the high level of beiging attained in response to cold. Furthermore, it assumes a functional interaction of lymphoid cells with activated MCS to induce complete beige remodeling. The differences in the origin of the immune cell populations involved in cold- versus Adrb3-induced beiging will be important to address in future studies looking into the immune implications in thermogenesis.

Our results further showed that the interferon/Stat1 signaling pathway is activated by CL suggesting an importance of these pathways in myeloid activation during CL-induced beiging. Previous work on human peripheral blood mononuclear cells (PBMC) showed that interferon synthesis was suppressed by catecholamines and favors a type 2 cytokine through Adrb2 stimulation (Wahle et al., 2005). Furthermore, neural inputs have been shown to increase lymphocyte numbers in vitro and in vivo (Agarwal and Marshall, 2000, Araujo et al., 2019). These studies along with our results support a model in which catecholamines released during cold exposure leads to lymphoid immune recruitment through the suppression of the interferon response activated by Adrb3 stimulation alone. Moreover, these results suggest the involvement of different coordinated signaling pathways to induce beige remodeling during cold and open a possibility that different signaling can lead to distinct adipocytes with equivalent beige phenotype.

In conclusion, these data provide a comprehensive atlas of the cellular dynamics during beige remodeling within white adipose tissue. We shed light on the complexity and the differences of both the immune and the transcriptional response of Adrb3- and cold-induced beiging. A better understanding of the signaling pathways and the cellular intra-organ communication influencing beige remodeling during Adrb3 and cold stimulation could ultimately lead to novel strategies to increase energy expenditure and protect against obesity.

## EXPERIMENTAL PROCEDURES

### Animals

C57Bl6 mice were purchased from The Jackson Laboratory at 6-week of age and acclimated for 2-week. Mice were housed in a temperature-controlled environment with a 12 hr light-dark cycle and ad libitum water and standard chow diet. For both single cells and flow cytometry experiments, 8-week-old mice were daily injected intraperitoneally (i.p.) with either vehicle (saline) or CL-316,243 (1 mg/kg) for 3 days before euthanasia. For cold exposure experiment, mice were maintained in 4°C room for 3 days. All animal studies were approved by the Boston University School of Medicine Institutional Animal Care and Use Committee.

### Histology

Tissue was fixed with paraformaldehyde, paraffin embedded, and sectioned (5 mm) prior to H&E staining or immunohistochemistry for Phospho-Stat1 (Tyr701) (58D6; cell signaling; 1:800).

### Immunofluorescence

5 μm slices of paraffin-embedded inguinal adipose tissue were mounted onto slides, deparaffinized and rehydrated before performing antigen retrieval. Tissue sections were stained with rabbit anti-CD3 (D7A6E; cell signaling, 1:200), rabbit anti-CD45 (D3F8Q; Cell Signaling, 1:100) overnight at 4C. After washing with 0.1% tween-20 TBS, sections were incubated for 1 hr at room temperature with fluorophore conjugated secondary antibody (donkey anti-rabit Alexa 647 (Invitrogen). Slides were then washed three times with 0.1% tween-20 TBS at room temperature in the dark. Coverslips were mounted using Prolong gold antifade (Thermofisher). Fluorescent images for all stained adipose tissue sections were captured with an Axio scan Z1 imager (Zeiss) at 20x magnification.

### Real-Time PCR

Total RNA was extracted from frozen tissues and cells using TRIzol reagent according to the manufacturer’s instructions. RNA concentrations were determined on NanaDrop spectrophotometer. Total RNA (100 ng to 1 mg) was transcribed to cDNA using Maxima cDNA synthesis (Thermo Fisher Scientific). Quantitative real-time PCR was performed on ABI Via detection system, and relative mRNA levels were calculated using comparative threshold cycle (CT) method. SYBR green primers are listed in Table S1.

### Flow cytometry analysis and data processing

Freshly isolated iWAT stromal vascular cell (SVC) were resuspended in FACS buffer (PBS/1% BSA). Samples were blocked with mouse Fc block (Biolegend; 1:50) for 5 min then incubated with antibody mix supplemented brilliant stain buffer (BD Biosciences) and monocyte blocker (Biolegend) for 20 min at 4°C protected from light. SVF cells suspension was rinsed 3 times before flow cytometry analysis with Aurora spectral cytometry analyzer (Cytek Biosciences). Antibodies are listed in Table S2. All data analyses was performed in Omiq.ai cloud cytometry data analysis platform. Single live CD45+ cells were clustered with Phenograph (Levine et al., 2015) and visualized with opt-SNE dimensionality reduction algorithm (Belkina et al., 2019). Groupings of clusters based on hierarchical clustering of median fluorescence intensities across multiple surface protein markers were annotated and color-overlaid on the opt-SNE projection of multidimensional data. Frequencies of each cell type were calculated from corresponding clusters and data were plotted and compared using Prism 6.0 (Graphpad).

### Isolation of Stromal Vascular Cells from Mouse iWAT

inguinal white adipose tissues (WAT) from control and CL- and cold-treated mice were collected after CL treatment and processed for SVC isolation using mouse adipose tissue dissociation kit (Miltenyi biotec) according to manufactures.

### Single Cell RNA Sequencing

Cells were prepared for single-cell sequencing according to the 10x Genomics protocols. Sequencing was performed on Illumina NextSeq500. The Cell Ranger Single-Cell Software Suite (v.3.1.0) (available at https://support.10xgenomics.com/single-cell-gene-expression/software/pipelines/latest/what-is-cell-ranger) was used to perform sample demultiplexing, barcode processing, single-cell 3’ counting, and counts alignment to mm10 mouse reference genome. For further analysis, the R (v.3.1) package Seurat was used (adapted workflow available at https://satijalab.org/seurat/v3.1/immune_alignment.html) (Stuart et al., 2019). Briefly, Cells with feature counts over 2500 or less than 200 or have over 5% of mitochondrial genes were filtered out. were filtered. All the samples were integrated and top 45 dimensions were used to generate the final clusters. Cells are represented with Uniform Manifold Approximation and Projection (UMAP) plots. The Seurat function “FindNeighbors” followed by the function “FindClusters” were used for clustering using resolution of 0.5. FindAllMarkers function was used to identify specific gene markers for each cluster. Violin plots were used to compare selected gene expression. Differential expression between clusters was obtained using MAST. Specific genes for each cluster were used for functional annotation and Go terms using Enrichr (Chen et al., 2013, Kuleshov et al., 2016).

### Statistical analysis

Data were analyzed using GraphPad Prism 6.0 software (GraphPad) and are presented as mean ± standard error of mean (SEM). Group comparisons were analyzed using either two-tailed unpaired student t test or a two-way ANOVA followed by multiple comparisons correction method stated in Figure legend. Differences were deemed statistically significant with p < 0.05.

### Code availability

Codes are publicly available in the relevant citations and custom script is available on request.

## ACKNOWLEDGMENTS

This work was supported by NIH/NIDDK grants DK117161 and DK117163. N.R was supported by American heart association (AHA) fellowship (17POST33660875). We thank Hu Tianmu and Yuriy Alekseyev of the BUSM Single Cell Sequencing Core for their advice and assistance. We also thank the Boston University School of Medicine (BUSM) Flow Cytometry Core Facility for support.

## AUTHOR CONTRIBUTIONS

Conceptualization, N.R. and S.R.F.; Methodology, N.R. and S.R.F.; Investigation, N.R.; Flow cytometry analysis N.R and A.C.B; Formal Analysis, N.R.; Mouse experiments, N.R, K.D, B.N.C; Writing – Review & Editing, N.R., and S.R.F.

---

**Supplemental Figure 1:**
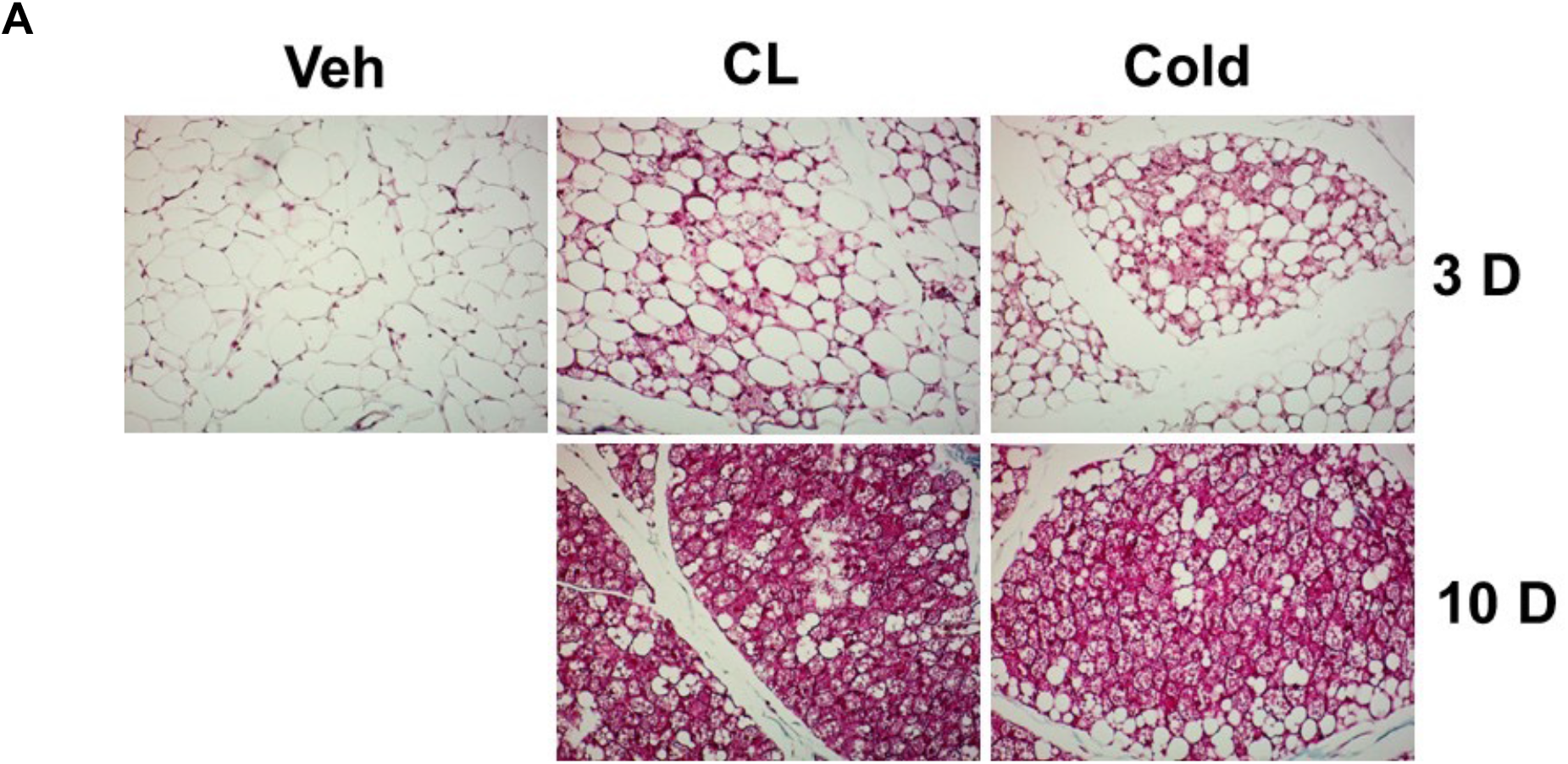
CL and cold treatment lead to distinct beige remodeling. (A) Representative H&E staining sections of from * weeks mice treated with vehicle, CL or cold for 3 and 10 days (n = 5–6 per group).

**Supplemental Figure 3:**
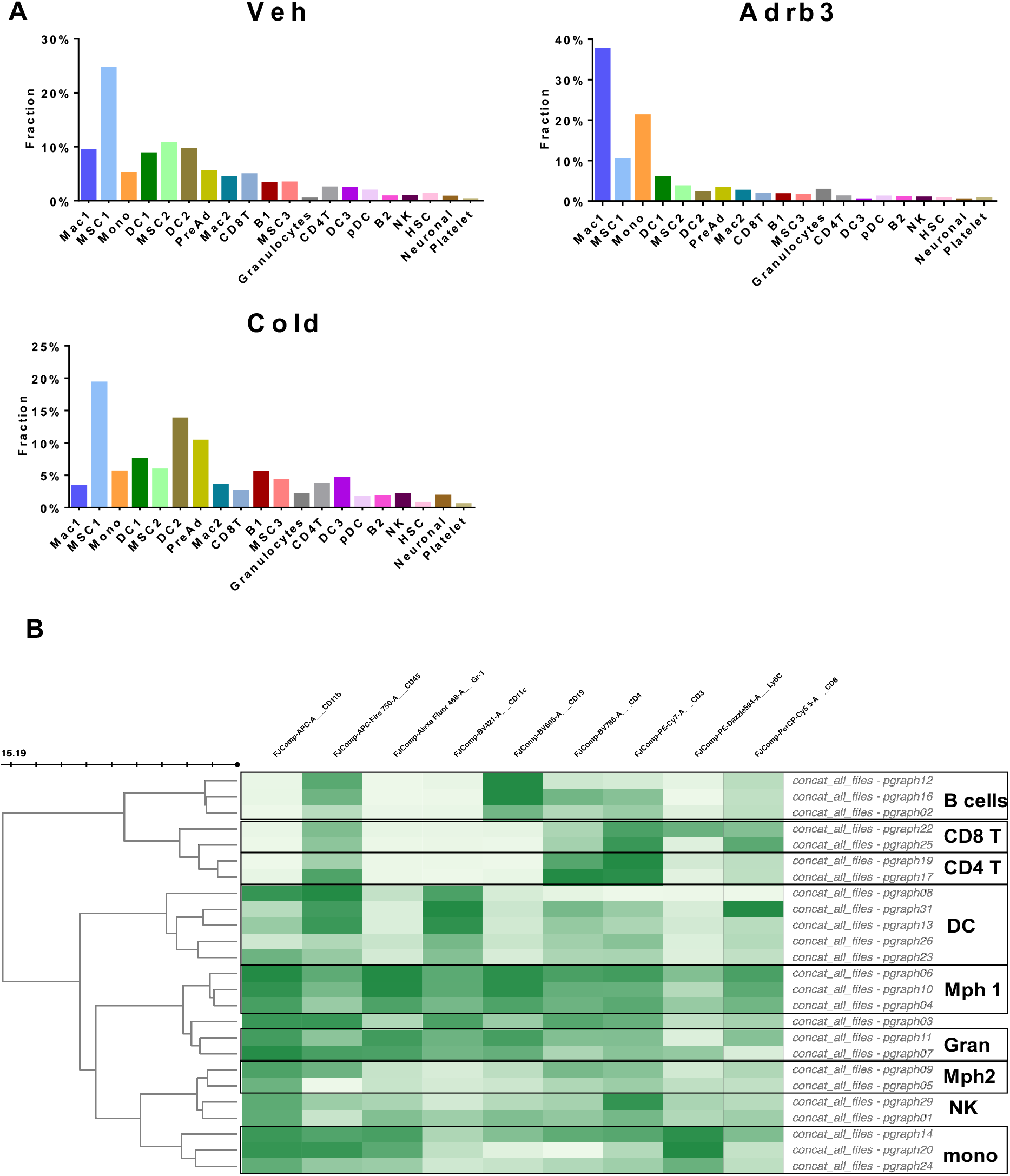
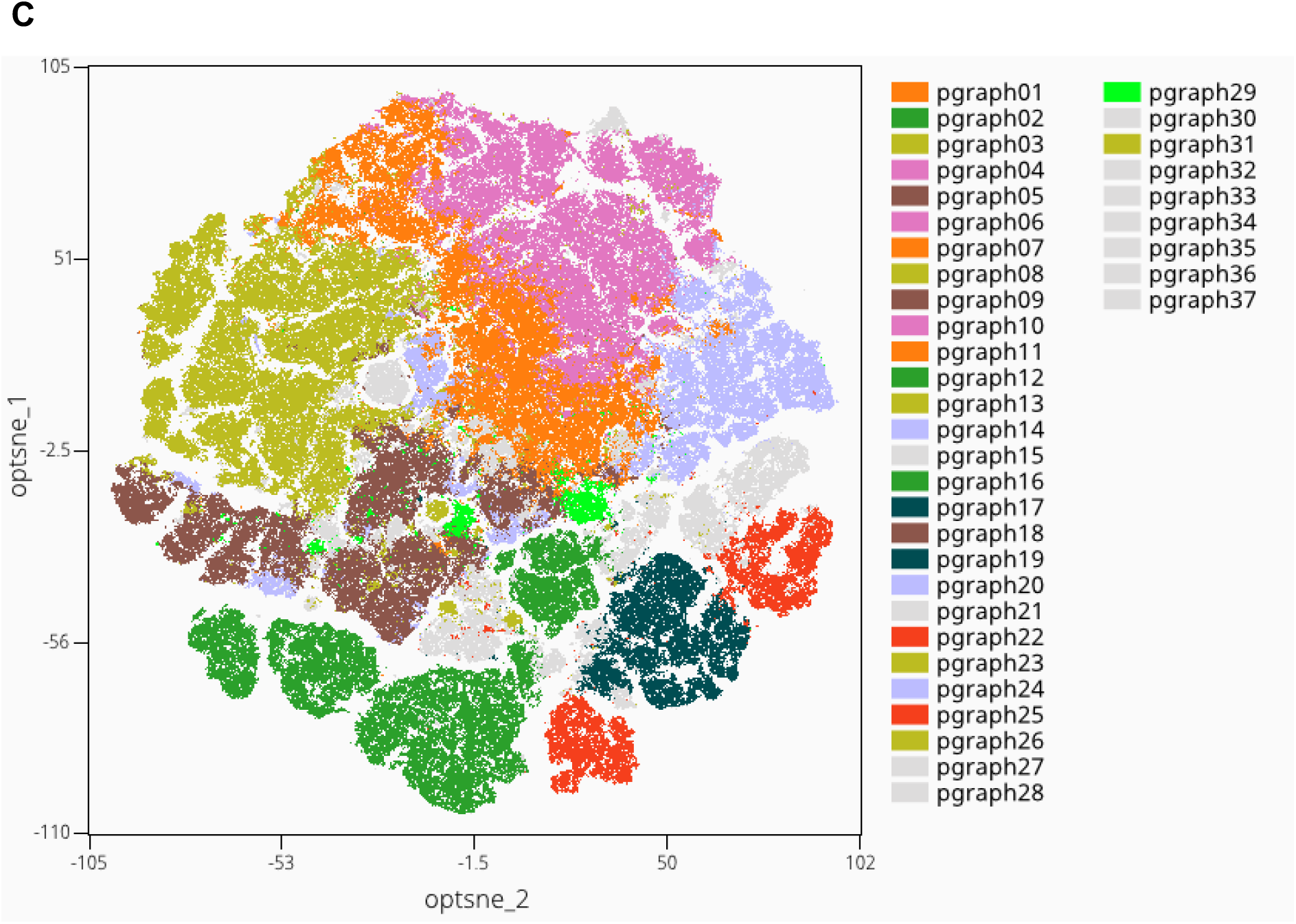
CL and cold differently alter the adipose resident immune compartment. (A) bar charts showing cells populations percentage per treatment from mice treated 3 days with either vehicle, CL or cold. (B) Heatmap of identified Phenograph clusters in flow cytometry data and their assignment to immune cells population of iWAT from mice treated 3 days with vehicle, CL or cold (each row represents a surface marker and Phenograph cell clusters and their assignment to immune cell types are shown in rows). (C) Flow cytometry opt-SNE plot of immune SVC in iWAT from mice treated 3 days with vehicle, CL or cold (All color overlays on t-SNE plots correspond to cell type classes; same color is assigned to Phenograph cluster groupings shown in panel B; grey color indicates debris clusters).

**Supplemental Figure 4:**
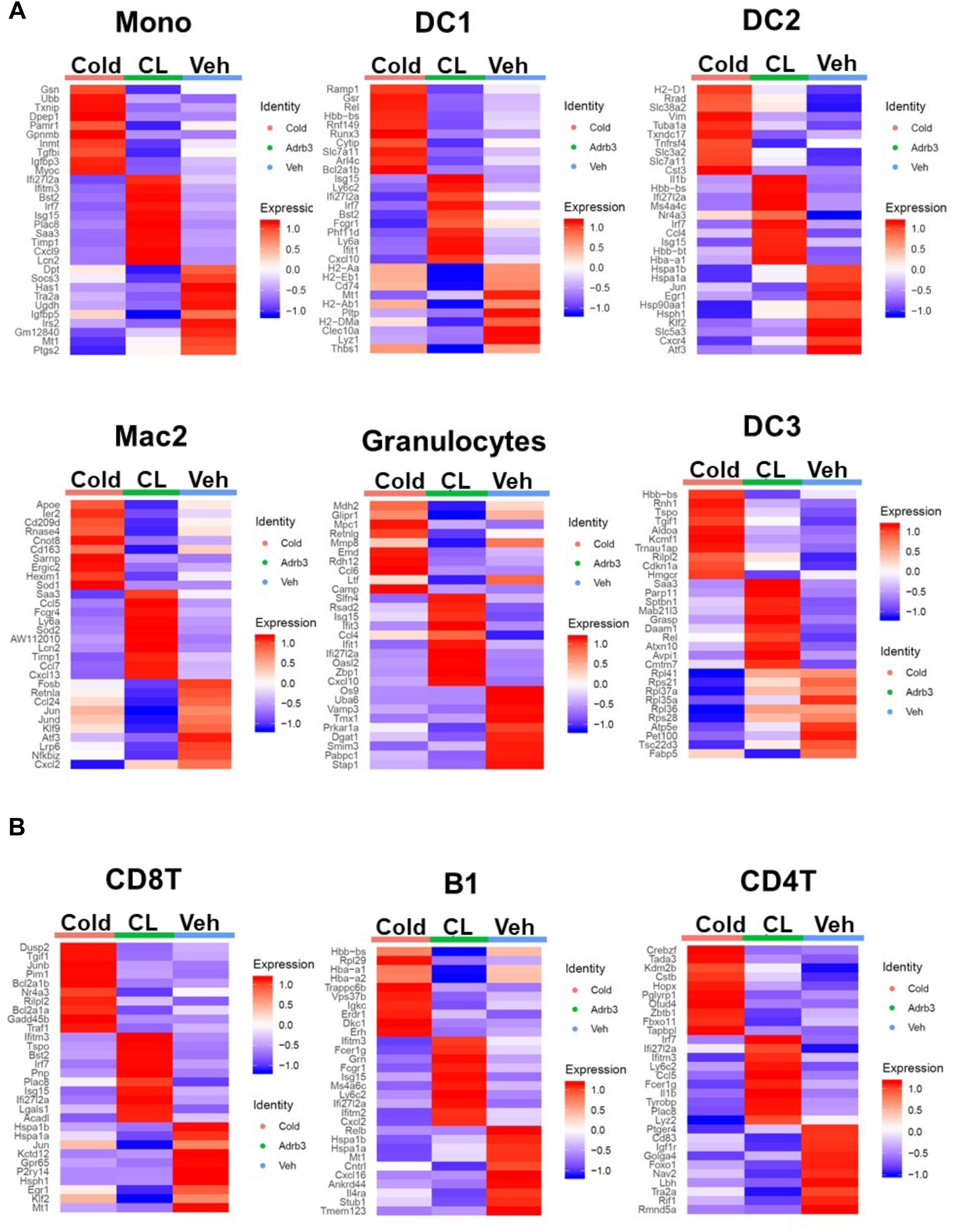
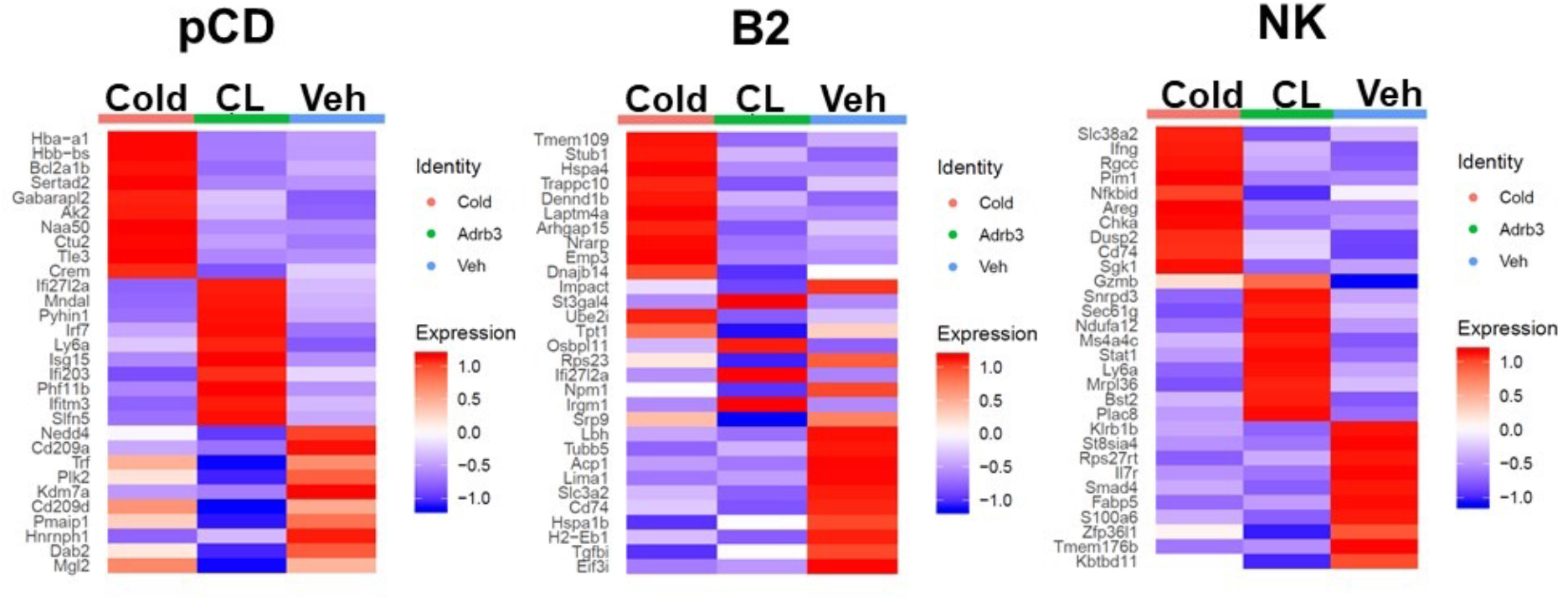
CL-induced beiging leads to an interferon/Stat1 response. (A) Heatmaps of differentially expressed genes in each myeloid cell type identified by single cell from mice treated 3 days with either vehicle, CL or cold. (B) Heatmaps of differentially expressed genes in each lymphoid cell type identified by single cell from mice treated 3 days with either vehicle, CL or cold.

## Inventory of Supplemental Information

### Supplemental Figures

Figure S1: Related to Figure 1. CL and cold treatment lead to distinct beige remodeling

Figure S3: Related to Figure 3. CL and cold differently alter the adipose resident immune compartment.

Figure S$: Related to Figure 4. CL-induced beiging leads to an interferon/Stat1 response.

### Supplemental Tables

**Table S1:**
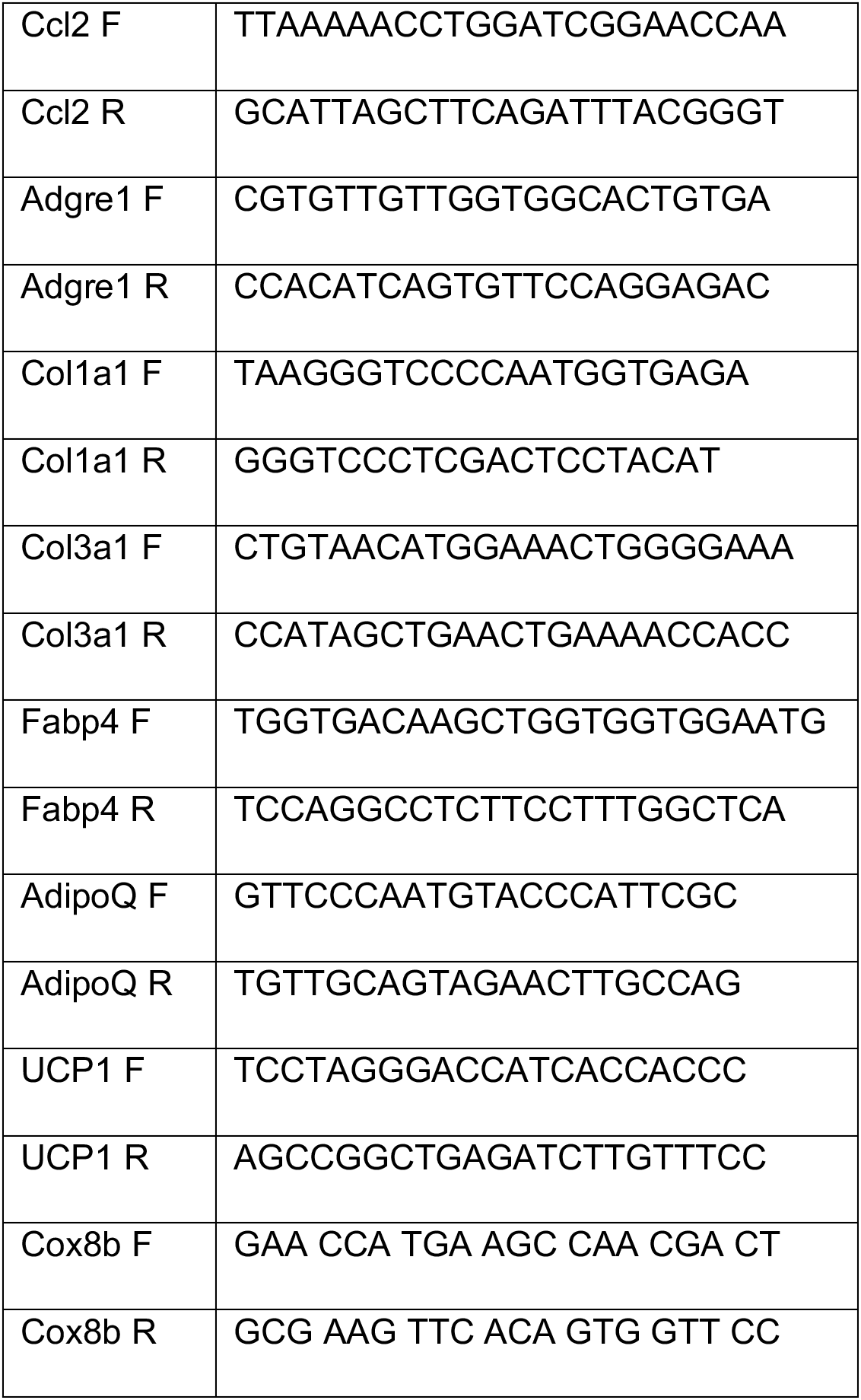
SYBR green primers list.

**Table S2:**
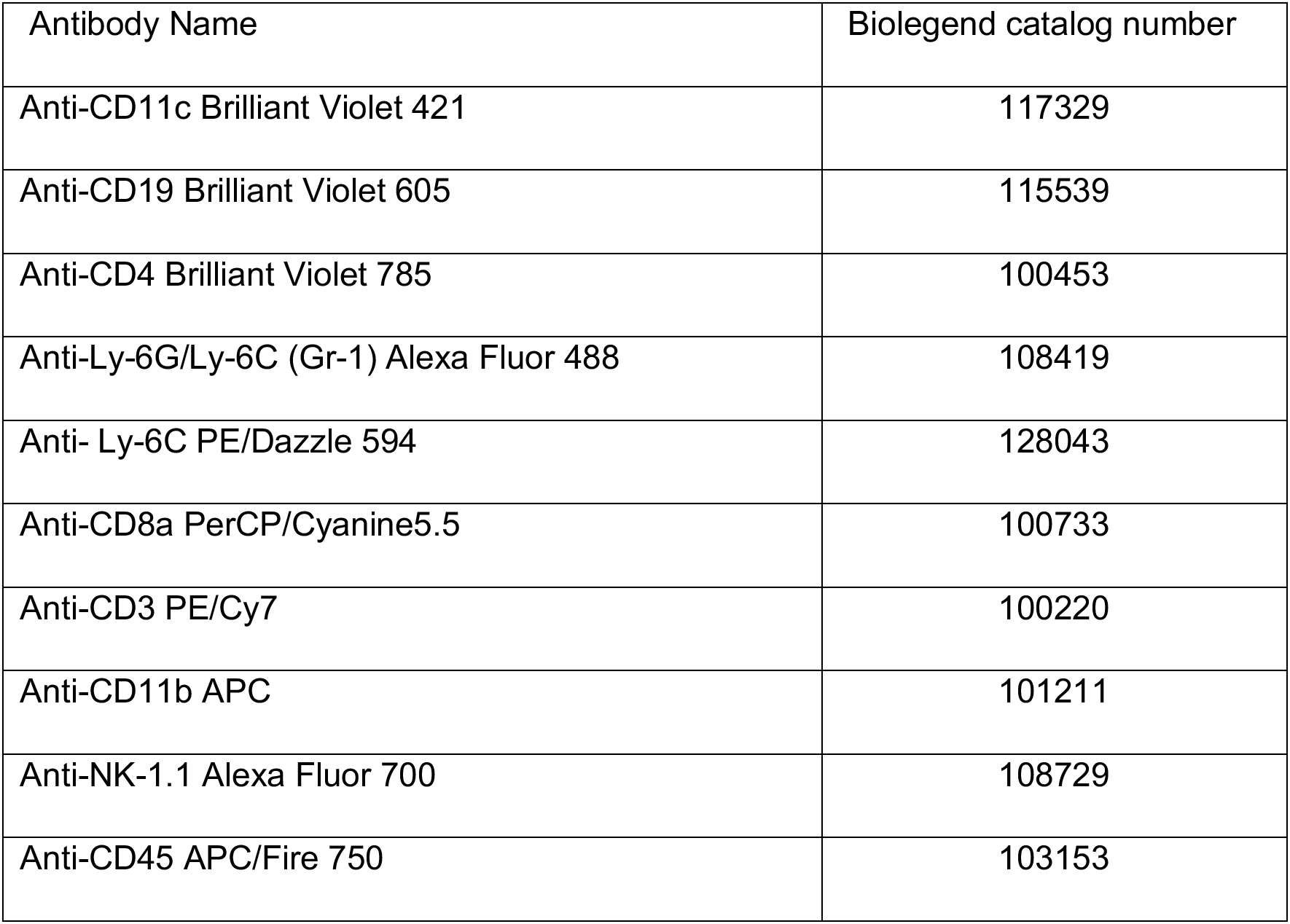
Flow cytometry antibodies list.

## REFERENCES

Agarwal, S.K., and Marshall, G.D. (2000). Beta-adrenergic modulation of human type-1/type-2 cytokine balance. J. Allergy Clin. Immunol. 105, 91–98.

Araujo, L.P., Maricato, J.T., Guereschi, M.G., Takenaka, M.C., Nascimento, V.M., de Melo, F.M., Quintana, F.J., Brum, P.C., and Basso, A.S. (2019). The Sympathetic Nervous System Mitigates CNS Autoimmunity via β2-Adrenergic Receptor Signaling in Immune Cells. Cell Rep. 28, 3120–3130.e5.

Bartelt, A., Bruns, O.T., Reimer, R., Hohenberg, H., Ittrich, H., Peldschus, K., Kaul, M.G., Tromsdorf, U.I., Weller, H., Waurisch, C., et al. (2011). Brown adipose tissue activity controls triglyceride clearance. Nat. Med. 17, 200–205.

Belkina, A.C., Ciccolella, C.O., Anno, R., Halpert, R., Spidlen, J., and Snyder-Cappione, J.E. (2019). Automated optimized parameters for T-distributed stochastic neighbor embedding improve visualization and analysis of large datasets. Nat. Commun. 10, 5415.

Berbée, J.F.P., Boon, M.R., Khedoe, P.P.S.J., Bartelt, A., Schlein, C., Worthmann, A., Kooijman, S., Hoeke, G., Mol, I.M., John, C., et al. (2015). Brown fat activation reduces hypercholesterolaemia and protects from atherosclerosis development. Nat. Commun. 6, 6356.

Burl, R.B., Ramseyer, V.D., Rondini, E.A., Pique-Regi, R., Lee, Y.-H., and Granneman, J.G. (2018). Deconstructing Adipogenesis Induced by β3-Adrenergic Receptor Activation with Single-Cell Expression Profiling. Cell Metab. 28, 300–309.e4.

Cannon, B., and Nedergaard, J. (2004). Brown adipose tissue: function and physiological significance. Physiol. Rev. 84, 277–359.

Chen, E.Y., Tan, C.M., Kou, Y., Duan, Q., Wang, Z., Meirelles, G.V., Clark, N.R., and Ma’ayan, A. (2013). Enrichr: interactive and collaborative HTML5 gene list enrichment analysis tool. BMC Bioinformatics 14, 128.

Choe, S.S., Huh, J.Y., Hwang, I.J., Kim, J.I., and Kim, J.B. (2016). Adipose Tissue Remodeling: Its Role in Energy Metabolism and Metabolic Disorders. Front. Endocrinol. 7, 30.

Cypess, A.M., Lehman, S., Williams, G., Tal, I., Rodman, D., Goldfine, A.B., Kuo, F.C., Palmer, E.L., Tseng, Y.-H., Doria, A., et al. (2009). Identification and importance of brown adipose tissue in adult humans. N. Engl. J. Med. 360, 1509–1517.

Eto, H., Suga, H., Matsumoto, D., Inoue, K., Aoi, N., Kato, H., Araki, J., and Yoshimura, K. (2009). Characterization of structure and cellular components of aspirated and excised adipose tissue. Plast. Reconstr. Surg. 124, 1087–1097.

Hepler, C., Shan, B., Zhang, Q., Henry, G.H., Shao, M., Vishvanath, L., Ghaben, A.L., Mobley, A.B., Strand, D., Hon, G.C., et al. (2018). Identification of functionally distinct fibro-inflammatory and adipogenic stromal subpopulations in visceral adipose tissue of adult mice. eLife 7.

Jojic, V., Shay, T., Sylvia, K., Zuk, O., Sun, X., Kang, J., Regev, A., Koller, D., Immunological Genome Project Consortium, Best, A.J., et al. (2013). Identification of transcriptional regulators in the mouse immune system. Nat. Immunol. 14, 633–643.

Kahn, C.R., Wang, G., and Lee, K.Y. (2019). Altered adipose tissue and adipocyte function in the pathogenesis of metabolic syndrome. J. Clin. Invest. 129, 3990–4000.

Kajimura, S., Spiegelman, B.M., and Seale, P. (2015). Brown and Beige Fat: Physiological Roles beyond Heat Generation. Cell Metab. 22, 546–559.

Kuleshov, M.V., Jones, M.R., Rouillard, A.D., Fernandez, N.F., Duan, Q., Wang, Z., Koplev, S., Jenkins, S.L., Jagodnik, K.M., Lachmann, A., et al. (2016). Enrichr: a comprehensive gene set enrichment analysis web server 2016 update. Nucleic Acids Res. 44, W90–97.

Lee, Y.-H., Kim, S.-N., Kwon, H.-J., Maddipati, K.R., and Granneman, J.G. (2016). Adipogenic role of alternatively activated macrophages in β-adrenergic remodeling of white adipose tissue. Am. J. Physiol. Regul. Integr. Comp. Physiol. 310, R55–65.

Lee, Y.-H., Kim, S.-N., Kwon, H.-J., and Granneman, J.G. (2017). Metabolic heterogeneity of activated beige/brite adipocytes in inguinal adipose tissue. Sci. Rep. 7, 39794.

Levine, J.H., Simonds, E.F., Bendall, S.C., Davis, K.L., Amir, E.D., Tadmor, M.D., Litvin, O., Fienberg, H.G., Jager, A., Zunder, E.R., et al. (2015). Data-Driven Phenotypic Dissection of AML Reveals Progenitor-like Cells that Correlate with Prognosis. Cell 162, 184–197.

Merrick, D., Sakers, A., Irgebay, Z., Okada, C., Calvert, C., Morley, M.P., Percec, I., and Seale, P. (2019). Identification of a mesenchymal progenitor cell hierarchy in adipose tissue. Science 364.

Nguyen, K.D., Qiu, Y., Cui, X., Goh, Y.P.S., Mwangi, J., David, T., Mukundan, L., Brombacher, F., Locksley, R.M., and Chawla, A. (2011). Alternatively activated macrophages produce catecholamines to sustain adaptive thermogenesis. Nature 480, 104–108.

Rajbhandari, P., Arneson, D., Hart, S.K., Ahn, I.S., Diamante, G., Santos, L.C., Zaghari, N., Feng, A.-C., Thomas, B.J., Vergnes, L., et al. (2019). Single cell analysis reveals immune cell-adipocyte crosstalk regulating the transcription of thermogenic adipocytes. eLife 8.

Ravasi, T., Suzuki, H., Cannistraci, C.V., Katayama, S., Bajic, V.B., Tan, K., Akalin, A., Schmeier, S., Kanamori-Katayama, M., Bertin, N., et al. (2010). An atlas of combinatorial transcriptional regulation in mouse and man. Cell 140, 744–752.

Sanchez-Gurmaches, J., and Guertin, D.A. (2014). Adipocyte lineages: tracing back the origins of fat. Biochim. Biophys. Acta 1842, 340–351.

Sharp, L.Z., Shinoda, K., Ohno, H., Scheel, D.W., Tomoda, E., Ruiz, L., Hu, H., Wang, L., Pavlova, Z., Gilsanz, V., et al. (2012). Human BAT possesses molecular signatures that resemble beige/brite cells. PloS One 7, e49452.

Stuart, T., Butler, A., Hoffman, P., Hafemeister, C., Papalexi, E., Mauck, W.M., Hao, Y., Stoeckius, M., Smibert, P., and Satija, R. (2019). Comprehensive Integration of Single-Cell Data. Cell 177, 1888–1902.e21.

Su, A.I., Wiltshire, T., Batalov, S., Lapp, H., Ching, K.A., Block, D., Zhang, J., Soden, R., Hayakawa, M., Kreiman, G., et al. (2004). A gene atlas of the mouse and human protein-encoding transcriptomes. Proc. Natl. Acad. Sci. U. S. A. 101, 6062–6067.

Sun, K., Kusminski, C.M., and Scherer, P.E. (2011). Adipose tissue remodeling and obesity. J. Clin. Invest. 121, 2094–2101.

Vishvanath, L., MacPherson, K.A., Hepler, C., Wang, Q.A., Shao, M., Spurgin, S.B., Wang, M.Y., Kusminski, C.M., Morley, T.S., and Gupta, R.K. (2016). Pdgfrβ+ Mural Preadipocytes Contribute to Adipocyte Hyperplasia Induced by High-Fat-Diet Feeding and Prolonged Cold Exposure in Adult Mice. Cell Metab. 23, 350–359.

Wahle, M., Neumann, R.P., Moritz, F., Krause, A., Buttgereit, F., and Baerwald, C.G.O. (2005). Beta2-adrenergic receptors mediate the differential effects of catecholamines on cytokine production of PBMC. J. Interferon Cytokine Res. Off. J. Int. Soc. Interferon Cytokine Res. 25, 384–394.

Yoshida, H., Lareau, C.A., Ramirez, R.N., Rose, S.A., Maier, B., Wroblewska, A., Desland, F., Chudnovskiy, A., Mortha, A., Dominguez, C., et al. (2019). The cis-Regulatory Atlas of the Mouse Immune System. Cell 176, 897–912.e20.

